# Broad distributions of sliding times are fingerprints of efficient target search on DNA

**DOI:** 10.64898/2026.03.21.713314

**Authors:** Jitin Rajoria, Arnab Pal

**Affiliations:** The Institute of Mathematical Sciences, CIT Campus, Taramani, Chennai 600113, India; Homi Bhabha National Institute, Training School Complex, Anushakti Nagar, Mumbai 400094, India

## Abstract

We investigate the target search process by proteins locating specific target sites along DNA – a phenomenon fundamental to biological functions such as gene regulation, transcription, replication, recombination, and gene-editing technologies. This process proceeds through a repetitive sequence of stochastic motions: consisting of one-dimensional (1D) sliding along the DNA contour interspersed with detachment and three-dimensional (3D) excursions in the bulk, and then reattachment to a random location on DNA. Recognizing this sequence of random events as analogous to the resetting processes widely studied in statistical physics, we employ a first-passage-renewal framework and derive general expressions for both the mean and fluctuations of the total search time. Our results are completely generic and do not depend on the detailed microscopic dynamics of either the 1D or 3D phases. Quite interestingly, we find that intermittent detachment can not only accelerate the mean search but can also regulate fluctuations around it. Our analysis reveals a universal fluctuation inequality that links the variability and mean of the sliding time to the mean excursion time, thereby identifying the fundamental conditions under which target search process becomes efficient. Notably, we find that broad distributions of sliding times emerge as a universal characteristic for optimal search efficiency—a feature emanating from the slow dynamics along the DNA. Using the facilitated diffusion mechanism as a representative example, we validate the generality of our results. These findings provide a unified theoretical framework connecting stochastic search, resetting dynamics, and biological efficiency, while also highlighting the crucial role of DNA structure such as its contour length in modulating search performance.

## Introduction

Biological processes involving target search on DNA, such as how transcription factors locate specific genes or how the CRISPR/Cas9 system identifies and eliminates foreign DNA in bacterial immunity, are fundamental to cellular functions.^1–4^ Transcription factors bind to specific regulatory regions on DNA, promoting or inhibiting the association of RNA polymerase and thereby modulating gene expression.^5^ Likewise, in the bacterial immune system, when viral DNA is integrated into the bacterial genome, the Cas9 protein, guided by RNA, searches for the complementary foreign sequence and cleaves it, ensuring genomic integrity.^4^ These cases exemplify protein–DNA recognition processes in which a protein must identify a functional binding site, typically only ~ 10–30 bp in size,^6,7^ among roughly 10^6^–10^9^ possible non-specific sites along the genome. Despite the small size of these targets relative to the entire DNA and the complexities posed by chromatin organization and molecular crowding,^8–10^ proteins locate their sites with remarkable speed. Understanding how such efficient and reliable search is achieved has long been a central question in molecular biophysics.

Extensive research has sought to elucidate how DNA-binding proteins (DBPs) efficiently translocate along DNA to locate their specific target sites. If the search relied solely on 3D diffusion in solution, the classical Smoluchowski rate for diffusion-limited reactions would be ~ 10^8^ *M*^−1^*s*^−1^.^11^ However, Riggs *et al*. demonstrated in *in vitro* experiments that the lac repressor locates its promoter even faster, indicating that 3D diffusion alone cannot account for the observed kinetics.^12^ Subsequent studies suggested that DBPs additionally scan DNA via one-dimensional (1D) diffusion,^13–16^ although extended sliding is limited by molecular crowding effects.^17^ A widely accepted and experimentally supported^18–21^ resolution is the facilitated diffusion mechanism, first conceptualized by von Hippel and Berg.^22–25^ In this strategy, a protein alternates between 1D sliding while bound to DNA and 3D excursions through the surrounding medium before re-binding at another location, see Figure 1. Facilitated diffusion has been observed both *in vivo* for the lac repressor^20,26–28^ and *in vitro*,^18,21^ and numerous studies have shown that this combined mode of motion dramatically accelerates the search process compared to purely 1D or 3D strategies.^29–41^

**Figure 1:**
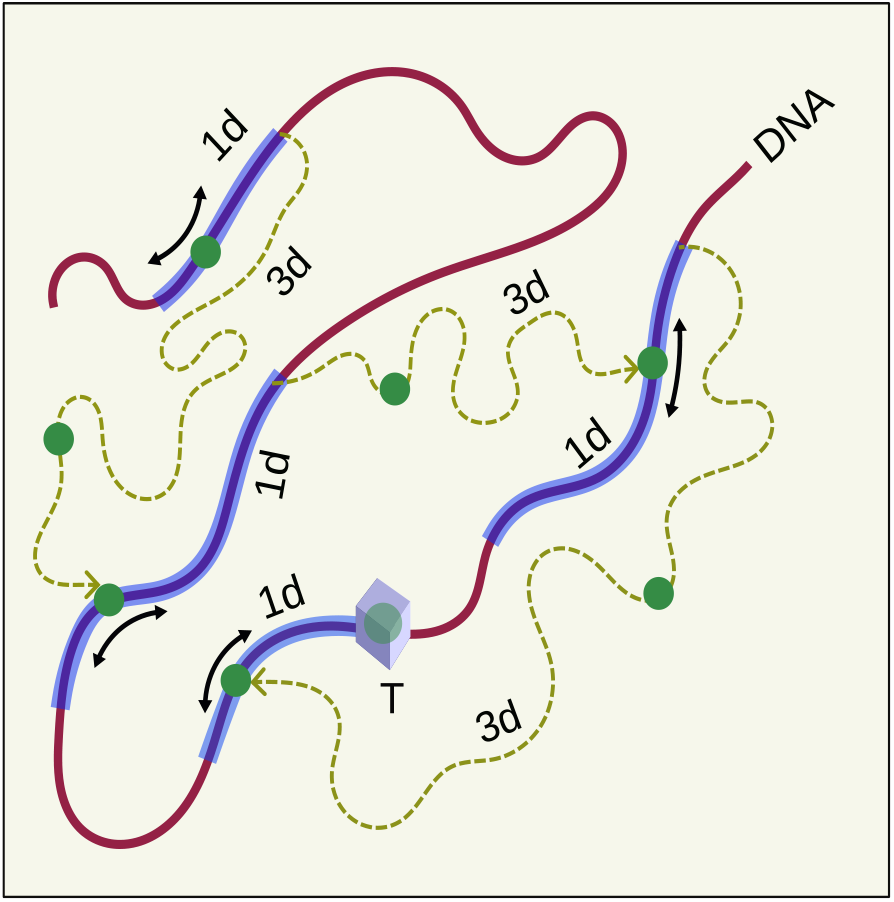
Schematic illustration of the target search process of a protein along the contour of a long DNA. The circle denotes the protein and the box represents the target site. The protein alternates between one-dimensional sliding along the DNA (solid line) and three-dimensional excursion in the cytoplasm (dashed line). After each 3D excursion, it reattaches to a random position on the DNA and resumes its search. This cycle repeats until the protein successfully locates the target. A circle–box pair indicates successful binding.

Although target search on DNA has been extensively investigated within the framework of facilitated diffusion, the classical picture of alternating 1D sliding and 3D excursions is not restricted to simple diffusive motion. In realistic cellular environments, the dynamics can be significantly altered by various physical and biochemical constraints. For example, the presence of DNA-bound obstacles^5,42–44^ and macromolecular crowding in the cytoplasm^5,8,17,45,46^ can substantially hinder protein mobility, leading in many situations to sub-diffusive behavior.^47–53^ These deviations from normal diffusion indicate that a theoretical description confined to standard Brownian motion may overlook essential aspects of the search dynamics. This naturally raises several key questions. Can one develop a unified formalism capable of describing the entire class of possible dynamical behaviors—diffusive, sub-diffusive, or otherwise—both along DNA and during 3D excursions? How do such altered dynamics influence the fluctuations in the total search time, beyond their impact on mean performance? Most importantly, is there a deeper physical principle that explains why intermittency—alternating between sliding and excursion phases—is such a successful strategy for molecular target search across biological systems? The goal of this work is to address these questions by establishing a general framework that captures the kinetics and stochastic features of facilitated target search on the DNA under minimal assumptions. By doing so, we aim to uncover the fundamental principles that enable intermittent search strategies to remain both robust and highly efficient within complex biological environments.

We develop a theoretical framework to describe the general stochastic process by which a DNA-binding protein searches for a target through alternating 1D sliding along DNA, detachment, 3D excursions, and subsequent reattachment. The search dynamics exhibit a natural renewal structure, akin to stochastic resetting, wherein a given system or a dynamics repeatedly restarts its exploration to complete a task^54–57^ or stochastically switches between the states or phases.^58,59^ Leveraging the first-passage under resetting formalism developed in,^60–62^ we obtain general expressions for both the mean search time and its fluctuations. A key strength of this approach is its independence from the microscopic details of the underlying dynamics—diffusive, anomalous, driven, or otherwise—making the conclusions broadly applicable. Within this unified framework, we uncover a universal fluctuation inequality that emerges as a hallmark of efficient target search. Strikingly, this relation reveals that for the search to be optimal, the distribution of 1D sliding times must be broad or heavy-tailed, implying that large fluctuations in sliding are not a drawback but rather a fundamental ingredient enabling efficient search by necessitating timely detachment and relocation. Furthermore, this inequality links the optimal detachment rate to properties of the DNA structure and associated kinetic parameters, providing a physical interpretation of when and why intermittent search becomes advantageous. As a concrete demonstration, we apply the theory to classical facilitated diffusion and show that its behavior is fully consistent with the universal principles identified here.

For brevity, we outline the structure of the paper below. Material and Method section introduces a general model for target search on DNA, delineates a renewal-based theoretical formalism, outlines the core assumptions followed by the expression for the mean total search time, emphasizing the generality of the approach. In there, we develop the universal *fluctuation inequality*, the key result of our analysis – given by (11), and examine its implications, particularly the role of broad time-distribution statistics in efficient target search mechanism. Moreover, we derive a general expression for the standard deviation of the total search time and provide the moment generating function, enabling computation of all higher moments. In the Results and Discussions section, the framework is applied to facilitated diffusion and validated against known results. This section further offers a quantitative analysis of inequality (11), including a phase diagram illustrating how facilitated diffusion depends on model parameters, identifying DNA length as a key control parameter and deriving the critical length. We also demonstrate a reduction in the search-time fluctuations due to the facilitated diffusion mechanism. Finally, in Conclusion section, we provide with a comprehensive summary of our findings and an outlook on future directions.

## Model setup and methods

### A generic model of target search on DNA

A generic target search process on DNA involves two different motions of protein: 1D sliding and 3D excursions. It can be further divided into three distinct kinetic phases (Fig. 1): sliding trajectory of the molecule (such as a protein) in 1D which terminates into detachment from the DNA before target localization; subsequent three-dimensional excursion of the protein in the surrounding medium followed by reattachment to the DNA; and the sliding of the protein along the DNA strand in 1D which eventually leads to the localization of target, and we call it a successful event. These events can be characterized by the random durations of detachment *T*_D_, 3D excursion *T*_3D_, and one successful event *T*_S_ in which target has been found (Fig. 2). In each attempt after binding on DNA, two outcomes are possible: (i) the protein locates the target with a characteristic time which we are denoting as *T*_S_ (a successful event), or (ii) it dissociates from the DNA after a time *T*_D_. In both cases, the protein remains in a sliding state along the DNA. Importantly, we do not assume the existence of a distinct chemical state in which the protein is passively “waiting” at a site to dissociate. Rather, the motion is continuously governed by sliding dynamics. The distinction between the two kinetic phases is therefore not based on different physical states, but on the outcome of a given search attempt: a “successful” phase corresponds to target localization prior to dissociation, while a “dissociation” phase corresponds to detachment before target finding. Thus, the dissociation phase is not treated as an independent physical process separate from sliding; instead, it represents the duration of a sliding trajectory that terminates in dissociation with a timescale *T*_D_. At this stage, we make no assumptions regarding the distributions of these times, which collectively determine the experimentally observable search or binding time(*T*_Tot_)—the time it takes a single molecule to locate the specific target site on DNA. We assume that the target can only be located through 1D sliding along the DNA, and not via the 3D excursions. The biological motivation for this assumption lies in the relative size of the target compared to the entire DNA. Typically, the target consists of only 10–30 base pairs^6,7^ among millions of base pairs present on the DNA. As a result, the fraction of the DNA corresponding to the target is extremely small, and the probability of directly locating the target via 3D diffusion from the cytoplasm is expected to be low. Therefore, most binding events suppose to occur at nonspecific sites, after which the protein locates the target through one-dimensional sliding combined with intermittent 3D excursions. This assumption has been followed in refs.^32,63^ as well. Figure 1 illustrates that the target is ultimately located by the protein during the sliding phase. We also assume that the target acts as a perfectly reactive site, meaning that as soon as the protein reaches the target, the reaction occurs instantaneously. This is valid when the target recognition process and binding to target are much faster than the 1D sliding. The target site is a high-affinity binding site. The initial conditions for every new start are random and uniformly distributed. We further assume that the 3D excursions are independent of the departure and arrival points. This restriction reflects the physical constraint that, during 3D diffusion, the protein typically re-associates with the DNA at random locations rather than specifically at the target site. Considering the two possible routes of search: uninterrupted 1D sliding or sliding-detachment-excursion-reattachment and renewal of search, the total random search time *T*_Tot_ can be calculated in the following way^34,60,62^

**Figure 2:**
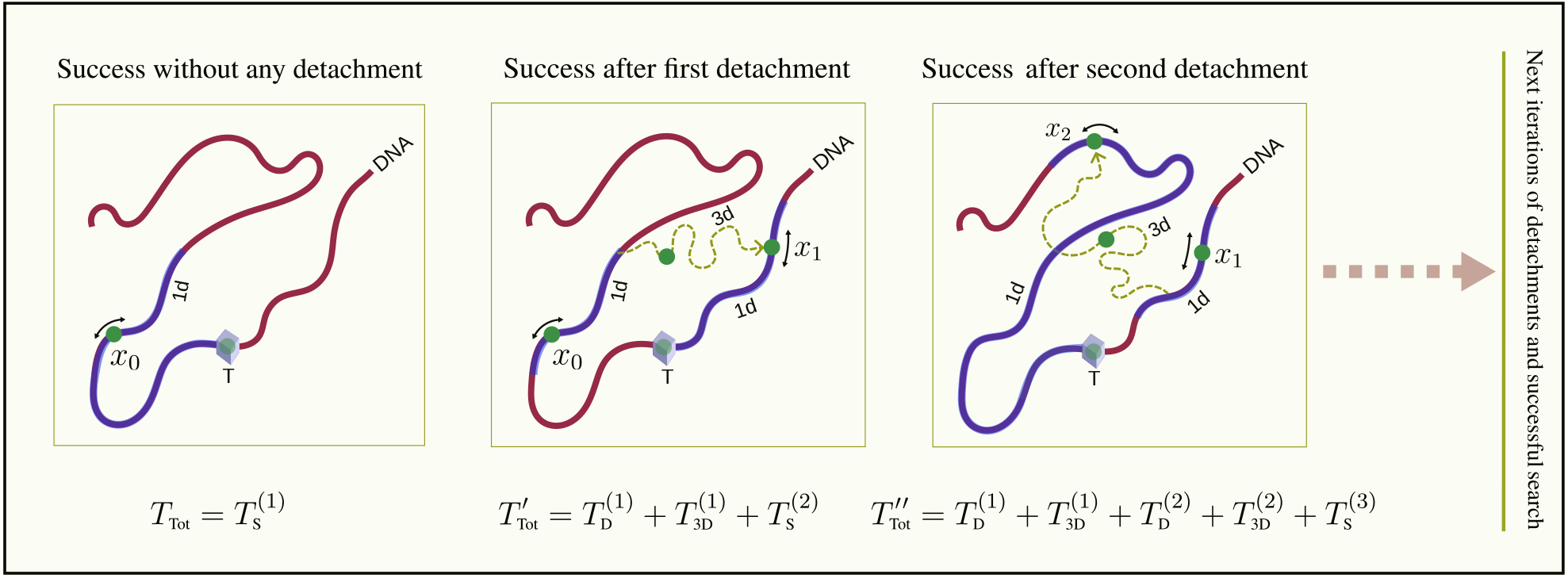
Schematic representation of the target search process as described by Eq. (1). The protein begins its search at position *x*_0_. If it finds the target without detaching from the DNA, the search is successful and ends (leftmost box), with total time 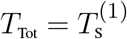. If the protein fails to find the target, it detaches after time 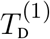, undergoes a 3D excursion of duration 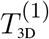, and rebinds at a new random position *x*_1_ on the DNA. A new search begins after the detachment. If the target is found in this round, the total search time 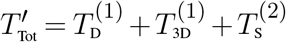 (middle box). In the rightmost box, we show another representative trial where the protein finds the target after two detachment events: it detaches after 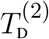, makes another 3D excursion of duration 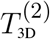, and binds at a new position *x*_2_, repeating the process. In this case, the total search time is 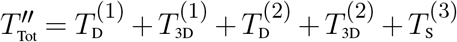. This is the repetitive renewal structure of the random search time *T*_Tot_ as used in Eq. (1), where 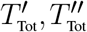, … are independent and identical copies of the total search time emanating from different search realizations. In the diagram, circles represent the protein, box denotes the target site, and a circle–box pair indicates successful binding.

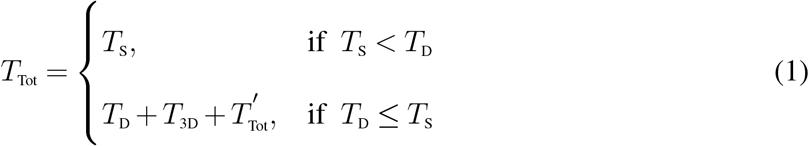

where 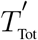 is is an identical and independent copy of *T*_Tot_. The above renewal equation can be interpreted in the following way: if the protein is able to find the target site without any detachment, the search therein ends (top row); otherwise the protein detaches from the DNA, completes a 3D excursion and reattaches to another location on DNA following which the search renews (bottom row). This series of stochastic events continues until the target is found, see Fig. 2. For the former event (a successful event), the timescales should satisfy the condition *T*_S_ *< T*_D_ which occurs with the following success probability

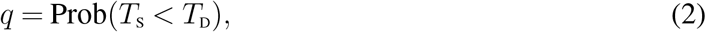

while the latter events occur with a complementary probability 1 − *q*. Note that *q* is often denoted as the committor in the chemical reactions.^64,65^ Notably, each renewal of the search process does not preserve any memory of the preceding search attempt. Throughout this work, we assume that the detachment rate and the ensuing 3D excursion–reattachment dynamics are independent of the protein’s position at the moment of detachment or reattachment. Situations where such spatial dependencies become relevant can still be incorporated by explicitly embedding them into the time scales *T*_D_ or *T*_3D_ within the renewal equation (1); the renewal structure remains valid as long as no memory is carried over between consecutive search trials.

### Mean search time

To move further, we rewrite Eq. (1) in the following way

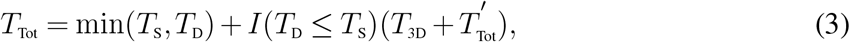

where the random variable min(*T*_S_, *T*_D_) is minimum of the two times *T*_S_ and *T*_D_, and *I*(*T*_D_ ≤ *T*_S_) is an indicator random variable that takes value unity when *T*_D_ ≤ *T*_S_ and zero otherwise. Taking expectation on both sides of Eq. (3) gives an exact expression for the average total search time

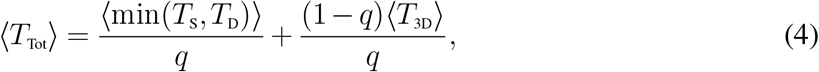

which is the first result of our analysis. Notably, this result does not depend on the specific nature of the individual kinetic phases and in principle, the timescales involved can be non-Markovian in nature. As mentioned earlier, the renewal structure pertains to the independence of successive search attempts and assumes that those are memory-less at the level of complete cycles. This assumption is justified by the fact that, upon detachment from the DNA and subsequent rebinding, the protein effectively loses memory of its previous trajectory, thereby initiating a new search attempt with identical statistical properties. This should not be confused with the underlying dynamics within each attempt, which can, in general, be non-Markovian. Importantly, the validity of this renewal structure does not require the dynamics within an individual search attempt to be Markovian. The motion during a single attempt may, in principle, exhibit memory effects, such as subdiffusive behavior. The renewal assumption only necessitates that such memory is reset between successive binding events, ensuring independence at the level of complete search cycles.

To move forward, we will assume that detachment is a Poisson process such that detachment times are exponentially distributed with a constant detachment rate (denoted by *λ*_1_), i.e., 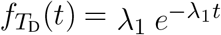. This assumption has been widely adopted in previous theoretical studies.^30,32,63,66,67^ Each detachment event of the protein from the DNA can be viewed as a barrier-crossing process that separates the bound (sliding) and unbound (3D diffusive) states.^30^ A closely related analogy exists in chemical kinetics, where the breaking of a molecular bond requires surmounting an activation barrier.^68,69^ The residence time in the bound state is characterized by an effective rate constant. By the same reasoning, protein–DNA detachment can be modeled as a thermally activated process. Consequently, the duration a protein remains bound to DNA before detachment is assumed to follow a single exponential distribution^30,70,71^ with rate *λ*_1_, reflecting the random and memory-less nature of such barrier-crossing events. However, our results can be generalized to non-exponential waiting time distributions, although in the present work we restrict ourselves to the exponential case for analytical tractability. More general non-exponential forms may arise due to heterogeneous binding landscapes, trapping in deep energy wells, conformational switching of the protein, or crowding effects. Such mechanisms can lead to memory-dependent detachment dynamics and broad distributions of dissociation times.

By taking an exponential distribution for the detachment time, we find

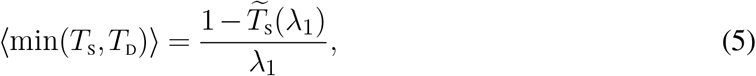

where 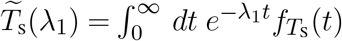 is the Laplace transform of the time distribution 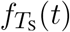 for the random variable *T*_S_ of a successful event.

Similarly, *q* can also be computed in the following way(see appendix - E and F)

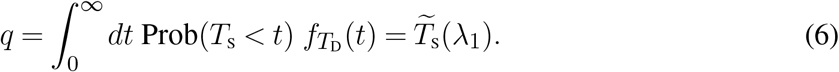

After substitution, we arrive at the following expression for the average total time to find the target on the DNA

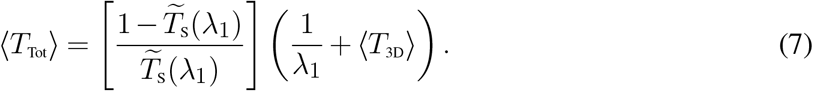

### Detachment events

The number of detachment events (denoted by 𝒩) till a successful search is also a random variable that can fluctuate between the realizations. This can also be measured by noting that a successful search is followed by *k*(*k* = 0, 1, 2, · · ·) unsuccessful detachment events and thus, the number should satisfy a geometric distribution, namely

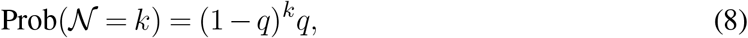

where *q* = Prob(*T*_S_ *< T*_D_) is the success probability. From this, we find the average number of detachment events to be

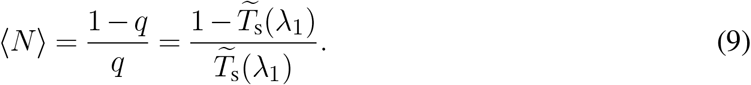

The key strength of the above expressions namely Eqs (4), (7), and (9) lies in its generality—it remains valid irrespective of the specific forms of the underlying 1D sliding dynamics of a successful attempt or 3D excursion processes. Moreover, the formulation naturally incorporates the Laplace transform which is analytically much easier to tract for a wide range of biologically relevant systems (see^63,72–74^ for a detailed review). For more complex scenarios, where analytical treatment becomes intractable, Eqs. (4) and (7) can be directly employed to numerically evaluate the mean search time using time-series data obtained from simulations or experiments. This makes the framework broadly applicable across both theoretical and data-driven investigations of search kinetics.

### A universal fluctuation inequality and the role of broad distribution statistics

The composite DNA-target search mechanism combining 1D sliding and intermittent 3D excursions have been demonstrated to be kinetically optimal, offering a compelling explanation for their prevalence in biological contexts.^67^ This suggests that living systems, through evolutionary refinement, have naturally evolved to exploit a synergistic interplay between 1D sliding along DNA and 3D excursions through the cellular milieu, thereby maximizing search efficiency. This observation prompts a more fundamental question: what underlying physical principle governs the enhanced efficiency of such coupled dynamics? In addressing this, we identify and derive a universal relation that encapsulates the intrinsic nature of these processes. Furthermore, we present several arguments supporting the notion that this relation is not a coincidental artifact, but rather a generic and robust feature underpinning such facilitated search processes in a broad class of biological systems.

In the combined search dynamics involving one-dimensional (1D) sliding along the DNA and intermittent three-dimensional (3D) excursions in the surrounding medium, an essential step for the transition between these modes is the detachment of the protein from the DNA strand. Drawing inspiration from this physical picture, we begin by analyzing the 1D sliding phase and introduce an infinitesimal small detachment, denoted by *δλ*_1_ for the protein to unbind from the DNA at any given instant. This small detachment rate effectively couples the 1D diffusion process to the subsequent 3D search in the bulk. For such a hybrid mechanism to achieve a net reduction in the overall mean search time compared to pure 1D diffusion, the gain from the 3D excursion must compensate for the time lost during detachment and rebinding. Consequently, the following condition must hold to ensure that the incorporation of the 3D phase indeed enhances the search efficiency:

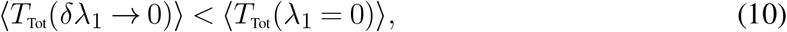

where ⟨*T*_Tot_(*λ*_1_ = 0)⟩ is the average time taken by a protein to locate its specific target site on the DNA when the detachment process (so the 3D excursion) is completely absent. In this case, the protein performs a purely one-dimensional (1D) sliding motion along the DNA strand without ever dissociating into the surrounding medium. Therefore, under this condition, ⟨*T*_Tot_(*λ*_1_ = 0)⟩ = ⟨*T*_S_⟩, the mean of a successful event in which target is located via uninterrupted 1D motion. Applying the inequality (10) in Eq. (7) and after a linear response calculation(see appendix -A), we arrive at the following relation

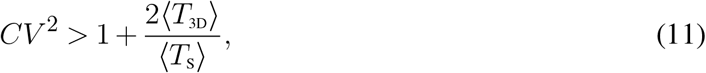

Where 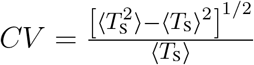, a ratio between the fluctuation and the mean of the time of a successful sliding event, is known as the coefficient of variation. Since this is a measure of statistical dispersion which estimates the fluctuations of a random variable around its mean, we call inequality (11) a *fluctuation inequality* or a *CV -criterion*. A larger coefficient of variation (*CV*) signifies a broader sliding time distribution, reflecting enhanced variability and stronger stochastic fluctuations, whereas a smaller *CV* corresponds to a narrower, more homogeneous distribution. Since the *CV* scales with the standard deviation, it naturally increases for systems exhibiting wide temporal dispersion.

Several comments are in order now. First, the inequality (11) tells us that for detachment to be beneficial, the *CV* for the underlying sliding time (*T*_S_) needs to be greater than unity plus a ratio involving the mean 3D excursion time (⟨*T*_3D_⟩), and the mean time of a successful sliding event(⟨*T*_S_⟩). Quite remarkably, the relation only relies only upon the mean and fluctuations of the sliding time and the 3D excursion time, and not on the higher order metric. Moreover, it remains robust to any underlying stochastic dynamics as long as these two measures are finite. Thus, the inequality is quite *universal* in nature.

Second, we would like to elucidate the implications of this fluctuation inequality in real physical scenarios. For instance, the protein dynamics along the DNA could significantly slow down due to the heterogeneous energy landscape along the DNA, where the presence of kinetic traps or sequence-dependent binding heterogeneities^30,31,75^ can transiently immobilize the protein, effectively slowing down the search process during the sliding phase.^29,76^ Moreover, the overall dynamics can become intrinsically slow due to the internal structural complexity of the protein. In particular, large or flexible macromolecules often exhibit anomalous diffusion, arising from conformational rearrangements, internal friction, or transient interactions with the environment. Such effects lead to sub-diffusive behavior, where the mean-squared displacement scales as ⟨*x*^2^(*t*)⟩ ~ *t*^*α*^ with 0 *< α <* 1 resulting in ultraslow relaxation kinetics.^48,77–80^ This intrinsic source of dynamic heterogeneity can substantially influence the sliding time. Consequently, both the mean sliding time and its fluctuations are enhanced, giving rise to multi-timescale or heavy-tailed sliding-time distributions, which may be well approximated by multi-exponential or power-law forms. These scenarios naturally satisfy the fluctuation inequality (11), since the left-hand side, characterized by a high *CV*, dominates as long as the 3D excursion times remain finite and relatively well-behaved.

In contrast, we also identify classes of protein–target binding mechanisms where detachment can only hinder the search, and we will show that these scenarios are also consistent with the constraints imposed by the fluctuation inequality (11). In one such simplest description, one could view the binding process as a sequence of Markovian transitions between *n*-kinetic states, each corresponding to a local minimum or a well on a coarse-grained energy landscape.^81–83^ In such cases, the distribution of the time for binding can be written in terms of the Erlang distribution 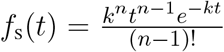 where *k* represents the equal rates at which transitions occur. In this case 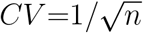 and no matter how the rates are rearranged, the fluctuations can not be reduced further. Naturally, such arrangement leads to a direct violation of the inequality nullifying the usefulness of detachment and 3D excursion in such target search problems. Another situation would be when the contribution from 3D excursions may become dominant in inequality (11), thereby violating the inequality. Physically, this situation arises when the 3D search dynamics are significantly slowed down due to macromolecular crowding in the surrounding medium. The cellular interior is densely packed with biomolecules such as water, ions, proteins, and enzymes,^45,46^ which generate both steric hindrance and viscoelastic effects. These factors restrict the diffusivity of the search protein during its excursions in the cytoplasm, leading to an increase in the average 3D search time ⟨*T*_3D_⟩ and hence altering the balance of timescales that determine the validity of the inequality. In essence, when the sliding-time distribution is narrowly peaked around its mean value ⟨*T*_S_⟩, one should have *CV <* 1 and thus, the inequality need not hold.

Overall, the balance between the timescales of the 1D sliding phase and the 3D excursions determines whether the system satisfies or violates the fluctuation inequality. This interplay underscores how subtle microscopic kinetic features—such as local trapping or diffusion heterogeneity—can give rise to pronounced macroscopic variations in search efficiency.

### Conditional times, distribution and fluctuations in search time

So far, we have only been concerned with the mean of the search time *T*_Tot_, but we will now move on to discuss the fluctuations and the full distribution by utilizing the renewal equation in Eq. (1). To this end, we will first define two stochastic conditional times as follows

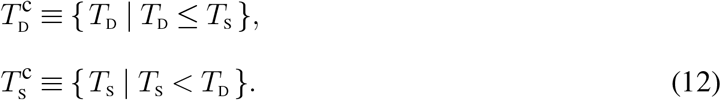

In words, 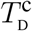 is the random detachment time conditioned on the fact that detachment is guaranteed to happen prior to the target location so that *T*_D_ ≤ *T*_S_ for all the realizations, see Fig. 3(a). Similarly, 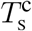 is the random time of a successful sliding given that the target has been found via sliding prior to detachment for all the realizations, refer to Fig. 3(b). The probability density of these conditional times can be written as^60^

**Figure 3:**
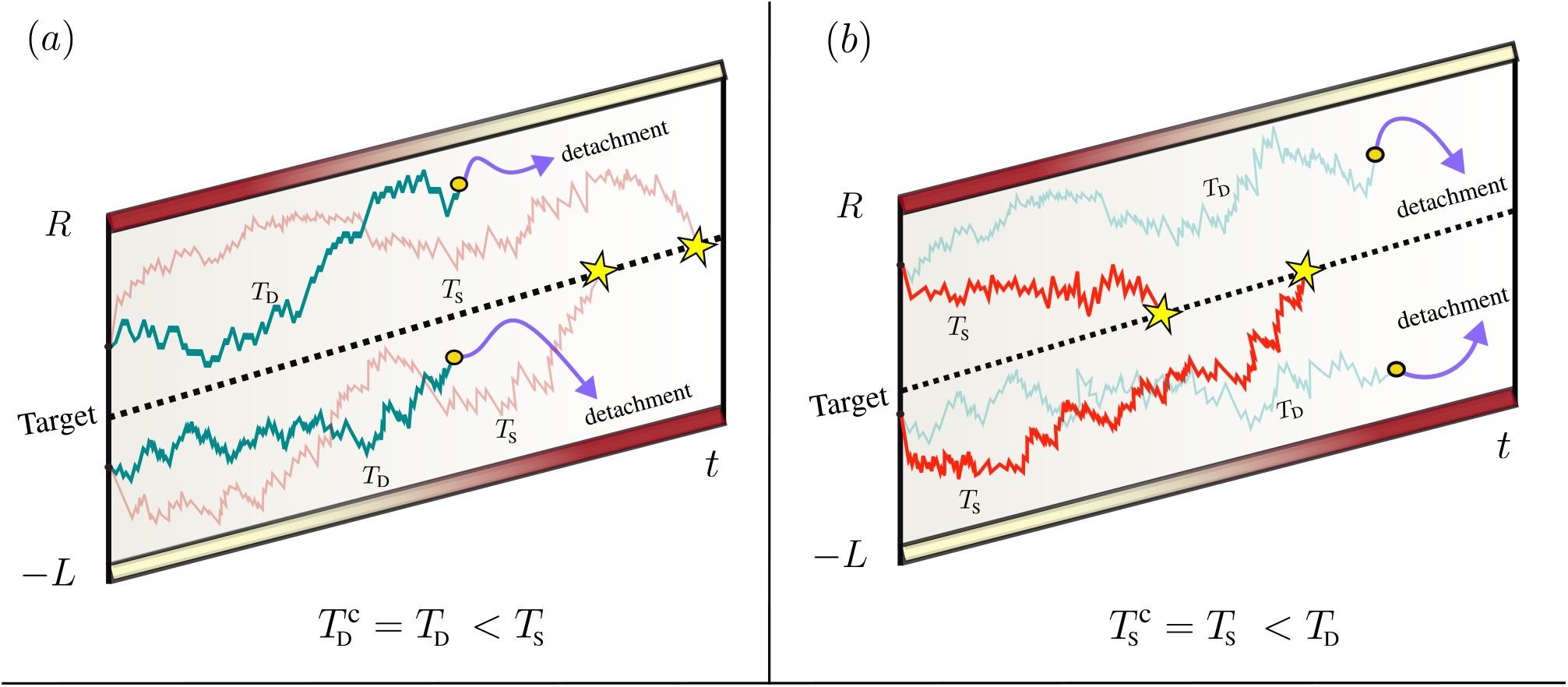
A schematic illustrating the stochastic conditional times. The figure depicts several possible protein trajectories in space–time as the protein moves within the domain bounded by *L* and *R*. Each path represents a distinct realization of the search process. The dashed vertical line indicates the target location, which remains fixed throughout the dynamics. Panel (a) illustrates the definition of the conditional time 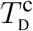. Two representative trajectories are shown starting from *t* = 0. In the first (green), the protein undergoes a detachment event at time *T*_D_, while in the second (light red), the protein locates the target (indicated by a yellow star) at time *T*_S_. In this example, *T*_D_ *< T*_S_, meaning detachment occurs before the target is found. Another realization (panel (a) below) satisfying the same condition, *T*_D_ *< T*_S_, is also depicted. By averaging over all such trajectories in which detachment precedes target encounter, one obtains the conditional time 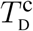. Similarly, panel (b) shows realizations in which the protein reaches the target before any detachment occurs. In these trajectories, the sliding event (red curve) leading to target encounter happens at time *T*_S_, while the corresponding sliding with subsequent detachment event (light green curve) would have occurred later, so that *T*_S_ *< T*_D_. Averaging over all such trajectories yields the conditional time 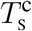, as defined in the main text. Together, the conditional times from panels (a) and (b), combined with their appropriate probability weights, form the basis for constructing the full distribution of the total search time.

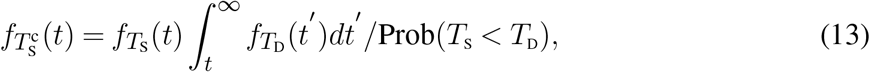

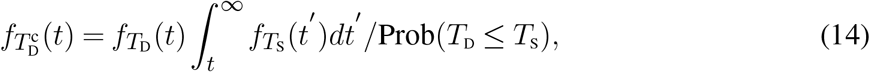

where *f*_S_(*t*) and *f*_D_(*t*) represent corresponding probability density functions of of the random variables *T*_S_ and *T*_D_ respectively.

Conditioning on these two possible set of events and applying the law of total expectations to the probability density function of the total search time in Laplace space, also known as the moment generating function, 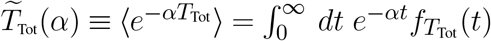, we arrive at the following expression for the distribution of the total search time in Laplace space

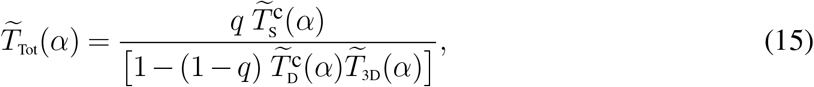

where *α* is a Laplace transform parameter. A formal derivation is given in appendix -C. As can be seen immediately, the above expression involves the Laplace transform of the conditional times, which need to be computed using Eq. (13) and Eq. (14).

Having the full probability distribution allows us to extract the mean and higher order moments of the total search time using the following relation

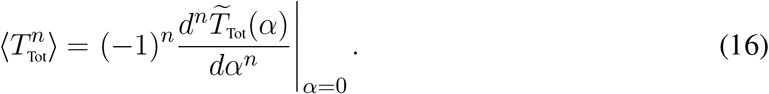

Differentiating once and setting *α*→0, we recover the mean total time as given by Eq. (4). Differentiating twice and setting *α*→0(see appendix-D), we obtain the second moment of the total search time

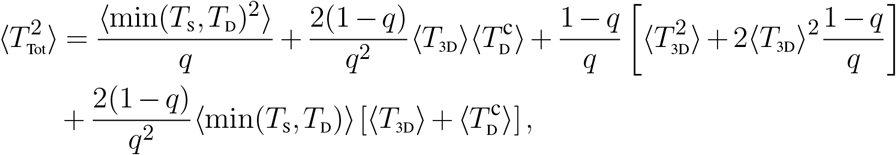

which is a general expression, independent of any particular nature of the sliding dynamics and 3D excursion. Moreover, we will be interested into the fluctuations for the search time, defined as

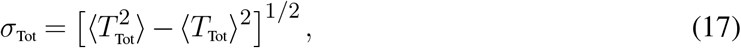

which will be studied in detail to further elucidate the role of detachment.

## Results and discussions

We will demonstrate in this section, by taking facilitated diffusion as a representative case, that the detachment process plays a dual role: it not only decreases the mean search time but can also optimally suppress the fluctuations around this mean, leading to a more reliable search process. In other words, controlled detachment can serve as an effective regulatory mechanism that balances speed and precision in target search. A detailed quantitative analysis of this optimization will be presented to further illustrate how the interplay between binding and detachment dynamics shapes the overall search efficiency.

### Facilitated diffusion as a target searching mechanism

So far, our analysis has remained general, without specifying the precise nature of the sliding motion along the DNA. We now focus on a more specific and biologically relevant framework—facilitated diffusion—a well-established mechanism by which DNA-binding proteins efficiently locate their specific target sites.^23–25,29–32,66,67^ In this mechanism, the search process alternates between two distinct modes: (i) 1D sliding of the protein along the DNA contour, and (ii) 3D excursions through the surrounding solution following detachment. In particular, the sliding phase is assumed to be a diffusive process, during which the protein moves along the DNA without detachment until it either finds the target or escapes at a stochastic detachment rate *λ*_1_. Upon detachment, the protein undergoes a 3D excursion, which we model as a rate process characterized by a constant reattachment rate *λ*_2_. While the 3D motion in a crowded cellular environment may, in reality, exhibit non-trivial or anomalous transport features due to macromolecular crowding,^8^ the rate-process assumption for the binding or association has been widely used in the literature and provides a reasonable first-order description.^31,32^

As we will demonstrate, this intermittent detachment–reattachment mechanism can play a crucial role in optimizing the overall search dynamics: it can significantly reduce both the mean search time and the associated fluctuations, transforming an otherwise inefficient search into a highly effective one. Finally, we emphasize that the simplifying assumptions used here do not affect our main conclusions—the key insight that intermittent dynamics optimizes search efficiency remains valid under fairly general conditions.

### Mean total time

In this framework, the DNA is modeled as a one-dimensional domain bounded by reflecting walls at *x* = −*L* and *x* = *R*, giving it an effective total length of *L* + *R*. The specific target site is located at *x* = 0 and treated as a point-like absorbing site, which can be detected only during the one-dimensional sliding motion of the protein. Direct binding to the target from three-dimensional excursions is not permitted. Once the protein encounters the target, the search process terminates.

The search begins from a random initial position *x*_0_ along the DNA. The protein slides along the DNA, searching for the target; if unsuccessful, it detaches, undergoes a three-dimensional excursion, and subsequently reattaches at a new random position to resume the search. The whole mechanism is illustrated in Fig. 4. This results in a sliding process with intermittent detachment and reattachment events—mechanistically analogous to stochastic resetting. Consequently, the analytical framework developed in the previous section can be directly applied to this system.

**Figure 4:**
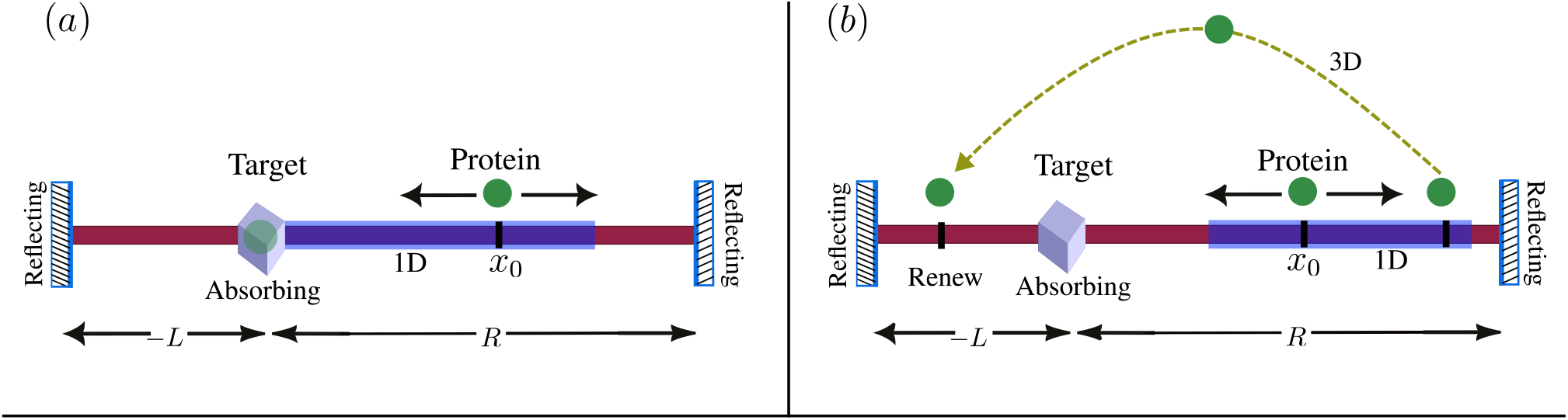
Facilitated diffusion model where a protein searches for a target site located on a one-dimensional DNA segment of total length *L* + *R*. The target site (box) is represented as an absorbing boundary, while the two ends of the DNA act as reflecting boundaries that confine the protein’s motion. Panel (a) indicates an event where the protein successfully detects the target while sliding. On the other hand, as shown on the panel (b), the protein can occasionally detach from the DNA after sliding on DNA, representing an unbinding event. Once detached, the search process effectively resets: the protein rebinds at a new, randomly chosen position along the DNA and resumes its search for the target. The shaded regions in both the panels indicate the extent of one dimensional sliding.

We begin by considering the motion of the protein along the DNA, which is assumed to be diffusive in nature. Accordingly, the time evolution of its position probability distribution function (PDF) can be expressed as^74,84^

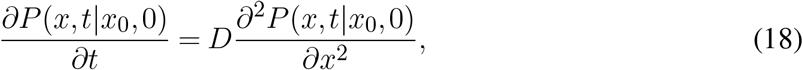

where *D* is the diffusion constant of one dimensional sliding motion and *P* (*x, t*|*x*_0_, 0) is the probability that the protein is at position *x* at time *t* given that it had started its search from the initial position *x*_0_ at time *t* = 0. Eq. (18) is supplemented with the following boundary conditions^63,74,84^

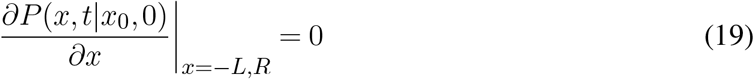

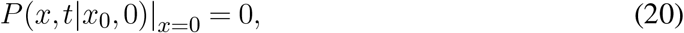

and initial condition *P* (*x, t*|*x*_0_, 0)|_*t*=0_ = *δ*(*x* − *x*_0_), where *x*_0_ is chosen uniformly along the DNA. The first set of boundary conditions implies that the DNA boundaries are reflecting, while the second one assumes that the target is an absorbing site. The latter condition reflects to the fact that the search ends as soon as the protein finds the target. The time distribution 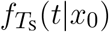 can easily be computed by employing the standard techniques as described in.^74,84^ To compute the mean search time using Eq. (7), we need to know 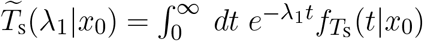 for which we can write the following backward equation (see appendix -B)

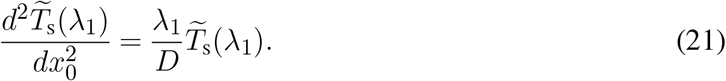

Following the appendix -B, we can write the solution as

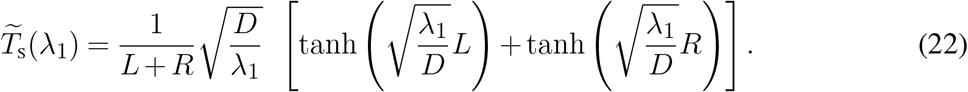

Substituting this expression in Eq. (7), we obtain the total mean search time^32,63^

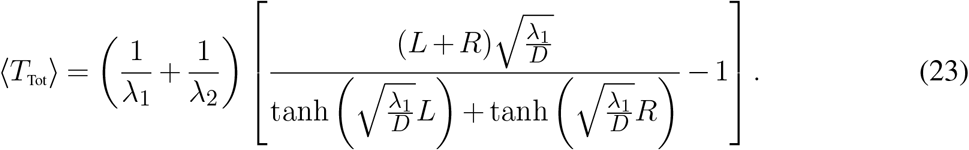

The average total time depends on various system parameters: the detachment rate *λ*_1_, attachment rate *λ*_2_, diffusion constant *D*, and the length of the DNA. In Figure 5(a), we present the variation of the mean total search time, ⟨*T*_Tot_⟩, as a function of the detachment rate. The DNA length is considered in units of base pairs. The different DNA lengths considered in Fig. 5(a) are 10^3^ bp, 1.5 × 10^4^ bp, 2.5 × 10^4^ bp and 5 × 10^4^ bp. The corresponding 3D excursions rates are 3 s^−1^, 15 s^−1^, 5 s^−1^ and 7 s^−1^ respectively. The diffusion constant is taken 5 × 10^5^ bp^2^/s, as measured in case of E. coli bacteria for *lac-repressor* protein.^16^ The values of *λ*_2_ and *D* are taken in a range similar to that reported in Refs.^32,63,66^ We observe that, for small values of *λ*_1_, the protein remains bound to local regions of the DNA for extended periods, exploring them slowly through one-dimensional diffusion. Owing to the inherently sluggish nature of diffusion, the search process in this regime becomes inefficient, leading to a large mean search time. Conversely, when *λ*_1_ is large, detachment occurs too frequently—the protein spends excessive time in the bulk, thereby reducing the overall search efficiency. Interestingly, for intermediate values of *λ*_1_, the mean search time decreases significantly, revealing the existence of an optimal detachment rate, denoted by 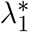 such that

**Figure 5:**
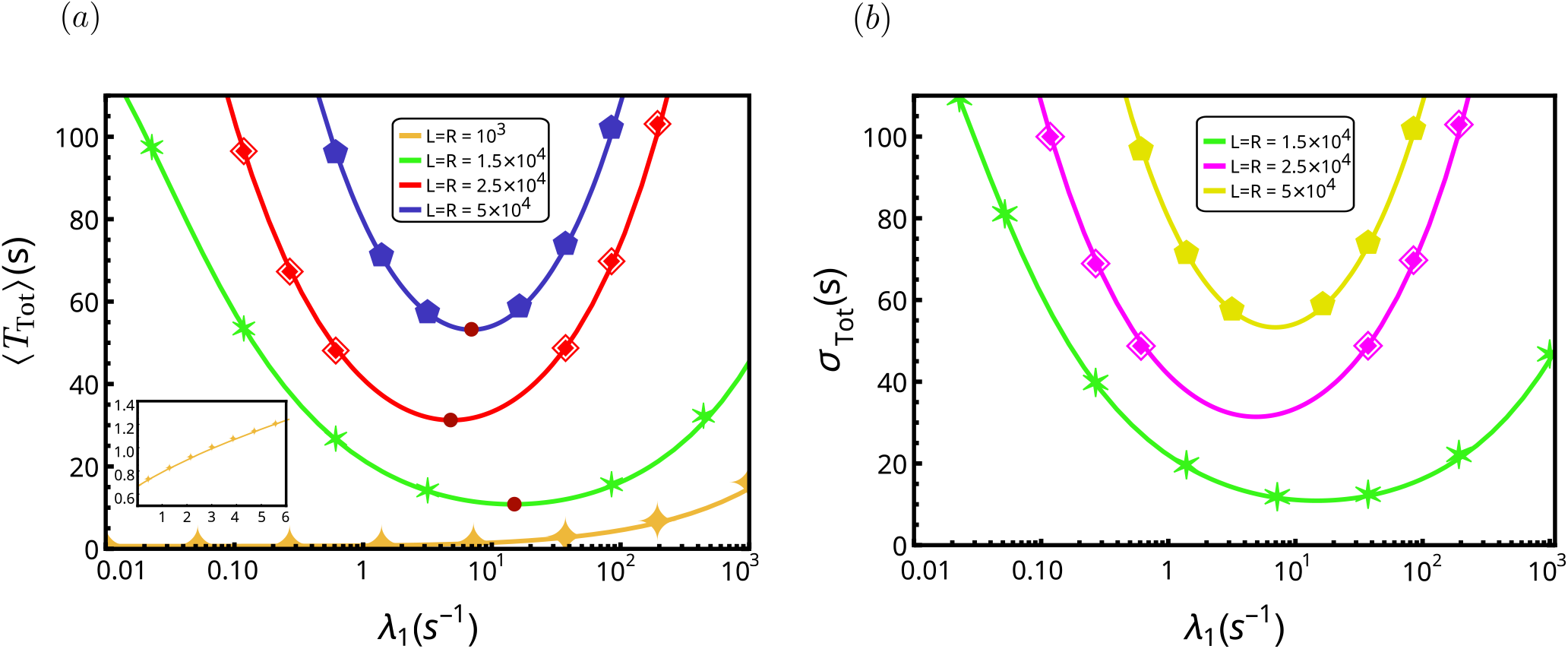
Mean and fluctuations of the total search time as functions of the detachment rate *λ*_1_ for a protein searching on a symmetric DNA. In both panels, solid lines denote analytical predictions and symbols represent simulation results. The diffusion constant is fixed at *D* = 5 × 10^5^ bp^2^/s, consistent with measurements for the *lac repressor* in *E. coli*.^16^ The reattachment rates are *λ*_2_ = 3, s^−1^ (monotonic case) and *λ*_2_ = 15, s^−1^, 5, s^−1^, 7, s^−1^ (non-monotonic cases). Panel (a) shows mean total search time ⟨*T*_Tot_⟩ versus *λ*_1_ for different contour lengths of the symmetric DNA. When present, an optimal detachment rate 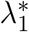 (circles) minimizes the mean search time. The inset corresponds to *L* = *R* = 10^3^, bp with *λ*_2_ = 3, s^−1^, where ⟨*T*_Tot_ ⟩ varies monotonically with *λ*_1_, indicating that detachment only slows the search and no optimum exists. This subtle yet important distinction between monotonic and non-monotonic behavior in the mean search time is precisely characterized by the fluctuation inequality given by inequality (11). Panel (b) shows standard deviation of the total search time *σ*_Tot_ as a function of *λ*_1_ for the same parameters. Introducing a finite detachment rate has a pronounced stabilizing effect, suppressing fluctuations in the search time by nearly 10^2^ orders of magnitude compared to the no-detachment case. This useful reduction highlights the essential role of detachment in regulating search dynamics.

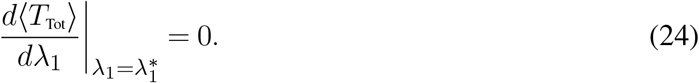

At this optimal point, the mean search time becomes the minimum. Physically, these detachment and reattachment events become optimally balanced such that the protein intermittently escapes the DNA and rebinds to distant sites, effectively bypassing redundant exploration and minimizing the overall search time.

In Fig. 5(a), we illustrate two distinct behaviors of the mean search time as a function of the detachment rate, *λ*_1_, for different DNA lengths. For certain DNA lengths, the curves exhibit a clear minimum, indicating the presence of an optimal detachment rate(marked) at which the mean search time is minimized. This optimality reflects the beneficial role of intermittent detachment, allowing the protein to efficiently escape local traps and re-initiate the search from new positions along the DNA. In contrast, for other DNA lengths, the curves display a purely monotonic increase in ⟨*T*_Tot_⟩ with *λ*_1_, implying that detachment only hinders the search and no optimal rate exists in this regime. This qualitative difference suggests that the DNA length fundamentally influences the balance between one-dimensional sliding and three-dimensional excursions. We argue that this transition between optimal and non-optimal search regimes can be understood within the framework of the universal fluctuation inequality discussed earlier, which provides a unifying explanation for how microscopic variability of the sliding times governs macroscopic search efficiency. This will be discussed next.

### Analysis of the *CV -criterion*

In this section, we show the universal fluctuation inequality (11) can be used to understand the optimization phenomena. To this end, we first compute *CV*^2^ for the 1D diffusion process

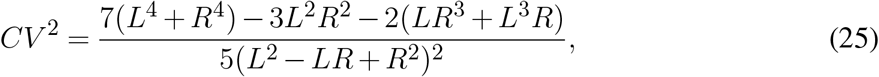

which is independent of the diffusion constant. For a symmetric DNA configuration (i.e., *L* = *R*), we obtain *CV*^2^ = 1.4, indicating that the relative fluctuation in the search time remains constant—independent of both the diffusion constant and the DNA length. In this scenario, the efficiency of the facilitated diffusion process is primarily dictated by the right-hand side of the *CV -criterion*, which involves the mean time of a successful sliding event, ⟨*T*_S_⟩, and the mean three-dimensional excursion time, ⟨*T*_3D_⟩. The latter is determined by the inverse of the reattachment rate, ⟨*T*_3D_⟩ = 1*/λ*_2_, while the mean sliding time during one-dimensional sliding is given by ⟨*T*_S_⟩ = *L*^2^/3*D*. Assuming that the diffusion coefficient *D* remains constant, increasing the DNA length leads to a quadratic growth in ⟨*T*_S_⟩, which in turn reduces the value of the right-hand side of the *CV -criterion*. Once the DNA length surpasses a certain threshold (to be computed below – see Eq. (26)), this term becomes smaller than the constant value of *CV*^2^, ensuring that the inequality is always satisfied. Beyond this threshold, facilitated diffusion operates in an efficient regime, where the balance between sliding and detachment yields an optimal search strategy.

We perform the analysis for Fig. 5(a) based on the *CV -criterion*. For symmetric DNA (*L* = *R*), the value of *CV*^2^ is fixed at 1.4. The right-hand side of the *CV -criterion*, however, varies for each DNA length and also the attachment rate. Table 1 summarizes the result of the analysis to illustrate these differences. For symmetric DNA length *L* = *R* = 1.5 × 10^4^ bp, 2.5 × 10^4^ bp, and 5 × 10^4^ bp, we find that *CV*^2^ *>* 1 + 2⟨*T*_3D_⟩/⟨*T*_S_⟩, see Table 1, and consequently the mean total time exhibits a minimum in Fig. 5(a). In contrast, for *L* = *R* = 10^3^ bp, *CV*^2^ *<* 1 + 2⟨*T*_3D_⟩/⟨*T*_S_⟩ and therefore no optimality in the mean total time is observed – see Fig. 5(a). To better understand the regimes in which facilitated diffusion becomes beneficial or detrimental, we present a comprehensive phase diagram in the parameter space spanned by (*λ*_2_, DNA length); see Fig. 6(a). The phase diagram clearly indicates two regions: facilitated diffusion beneficial, facilitated diffusion detrimental. The separatrix between the two regions can be directly obtained from the *CV -criterion* (11) by setting *CV*^2^ = 1 + 2⟨*T*_3D_⟩/⟨*T*_S_⟩ in the phase space of (*λ*_2_, DNA length) such that each point along the locus satisfies the above equality. Although demonstrated for the symmetric DNA, the fluctuation inequality more generally captures how the interplay between sliding and detachment governs an intrinsic optimization principle of facilitated diffusion and furthermore, constructing a practical phase diagram in the relevant parameter space.

**Table 1:** Table showcasing numerical values related to the *CV -criterion* from inequality (11). Parameters used: attachment rate *λ*_2_ = 3 s^−1^, 15 s^−1^, 5 s^−1^ and 7 s^−1^,^32,63^ diffusion coefficient *D* = 5 × 10^5^ bp^2^/s^16,32,63^ as observed in case of E. coli bacteria for *lac-repressor* protein;^16^ the coefficient of variation *CV*^2^ = 1.4 calculated when the length is symmetric around the target, i.e. *L* = *R* = length. While the first row fails to satisfy the *CV -criterion*, the remaining three fulfill it, thereby identifying—through the DNA contour length—the regimes in which the target search is efficient.

| length (bp $\times 10^3$) | $\langle T_s \rangle$ (s) | $\lambda_2$ ($\text{s}^{-1}$) | $\langle T_{3D} \rangle$ (s) | $1 + 2\langle T_{3D} \rangle / \langle T_s \rangle$ |
| --- | --- | --- | --- | --- |
| 1 | 0.66 | 3 | 0.33 | 2 |
| 15 | 150 | 15 | 0.06 | 1.0008 |
| 25 | 416 | 5 | 0.2 | 1.0009 |
| 50 | 1666 | 7 | 0.14 | 1.0001 |

**Figure 6:**
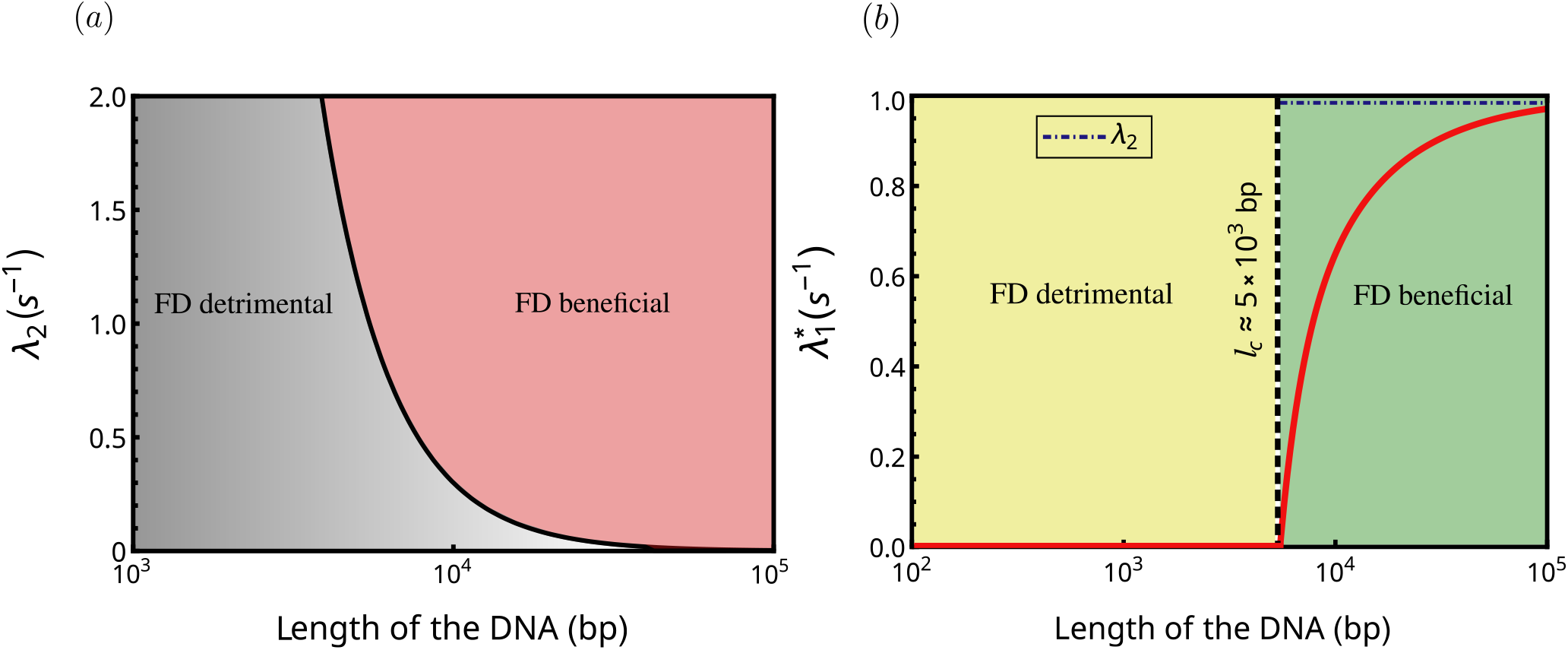
Panel (a) shows the phase diagram generated from the fluctuation inequality (11) in the *λ*_2_–DNA length parameter space. The diagram delineates two distinct regimes: one in which facilitated diffusion enhances the search efficiency and another in which it fails to provide any acceleration. The boundary (solid black curve) separating these regions—the separatrix—is determined by the equality *CV*^2^ = 1 + 2 ⟨*T*_3D_ ⟩/ ⟨*T*_S_ ⟩, which follows directly from the fluctuation inequality (11). Parameter combinations above this curve correspond to efficient search dynamics with an optimal detachment rate, whereas those below the separatrix yield monotonically increasing search times with no optimality. Panel (b) shows the plot of the optimal detachment rate 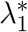 as a function of the DNA contour length. A finite, non-zero value of 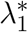 signifies that intermittent dynamics—i.e., facilitated diffusion—offers a genuine advantage over pure 1D sliding. The dashed vertical line marks the critical DNA length *l*_*c*_, obtained from the fluctuation inequality (11), beyond which detachment becomes beneficial and an optimal rate emerges. Below this threshold, the search time grows monotonically with *λ*_1_, and facilitated diffusion provides no improvement. The horizontal dot–dashed line highlights that for sufficiently long DNA, 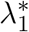 approaches the reattachment rate *λ*_2_ = 1 s^−1^, reflecting an effective balancing between the 1D sliding and 3D excursion phases at optimality. The diffusion constant in both the plots is fixed at 5 × 10^5^ bp^2^/s, as reported for the *lac-repressor* protein in *E. coli*.^16^

### Variation of optimal detachment rate with length of the DNA

From the preceding discussion, we have identified the length of the DNA to be a key parameter to understand the efficiency of the target search process. To strengthen the discussion further, here we analyze the behavior of the optimal detachment rate 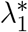 with respect to the total length, denoted by *l* = 2*L*, for the symmetric DNA. Analyzing a transcendental equation emanating from Eq. (24), we plot 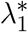 for different DNA lengths in Figure 6(b). By setting the diffusion constant to 5 × 10^5^ bp^2^ s^−1^, as reported for the *lac-repressor* protein in *E. coli*,^16^ and the attachment rate to 1 s^−1^, we observe that 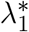 has a finite value for certain ranges of DNA length reestablishing the fact that facilitated diffusion is indeed useful. However, as the DNA length decreases, the value of 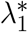 also decreases and eventually becomes zero at a critical DNA length. We denote this critical length of DNA by *l*_*c*_. If the DNA length decreases further below *l*_*c*_, then 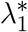 remains fixed to zero i.e., detachment hinders the target search.

We find numerically that the optimal detachment rate is zero for DNA length *l*_*c*_ ≈ 5 × 10^3^ bp. This critical length of the DNA, for facilitated diffusion to remain effective, can also be derived by substituting the expressions for *CV*^2^ from Eq. (25) and the mean time of a successful sliding event and 3D excursion time in the *CV -criterion*, yielding

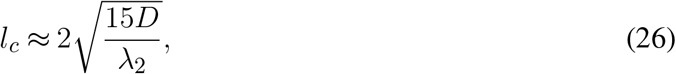

where we use the ‘≈’ symbol to emphasize that this critical value may vary in real systems where additional parameters also contribute. In addition, Fig. 6(b) indicates that as the DNA length becomes large, the optimal detachment rate 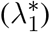 asymptotically approaches the attachment rate (*λ*_2_), which is taken as 1 s^−1^. Therefore, for sufficiently long DNA, the mean total search time ⟨*T*_Tot_⟩ attains its minimum value at 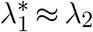. This observation was first noted by Slutsky and Mirny in^31^ and was later argued technically by Coppey *et al*.^32^ In summary, we show that the emergence of both the critical contour length and the optimal detachment rate are intimately connected, arising naturally from the *CV -criterion*, which constitutes the central and most significant result of this work.

### Role of 3D excursion timescales from^30,67^ and a prediction towards an efficient search

This section examines the role of three-dimensional (3D) excursion times in determining the efficiency of the search process. In the previous section, the rates of 3D excursions typically have values lying in the range 1 to 15 s^−1^, which are inspired from the Refs.^32,63,66^ In this section, we analyze an alternative set of parameters with ⟨*T*_3D_⟩ = 1*/λ*_2_. The estimated values of 3D excursions time spent by protein in cytoplasm are found to be vary from few milliseconds to hundreds of milliseconds.^30,85^ We again show a phase space diagram in parameter space of *λ*_2_ and the length of the DNA. In Fig. 7, we can easily notice that the region, in which facilitated diffusion is beneficial, has increased when the phase diagram is plotted for the higher values of *λ*_2_ (order of 10^3^ s^−1^). Recall that, in our earlier analysis, the length of the DNA emerged as a key parameter that governs the nature of the search mechanism. We therefore revisit the phase diagram to illustrate the effect of increased three-dimensional (3D) excursion rates for various DNA length. The results show that facilitated diffusion (FD) remains an efficient search mechanism even for short DNA lengths, provided that 3D excursions are also exceptionally fast. To understand clearly, we again extract the values of optimal detachment rate for different lengths of the DNA and plot it, see Fig. 7(red curve). We have considered that the average 3D time (⟨*T*_3D_⟩) for this particular curve is 10^−4^ s^30,67^ resulting in *λ*_2_ ~ 10^4^ s^−1^. It is observed that the optimal detachment rate 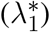 exists even for small value of DNA length. The critical DNA length (*l*_c_) is very short, i.e., approx 50 bp. The presence of a critical DNA length does not compromise the efficiency of the search mechanism, and the *CV -criterion* remains valid always.

**Figure 7:**
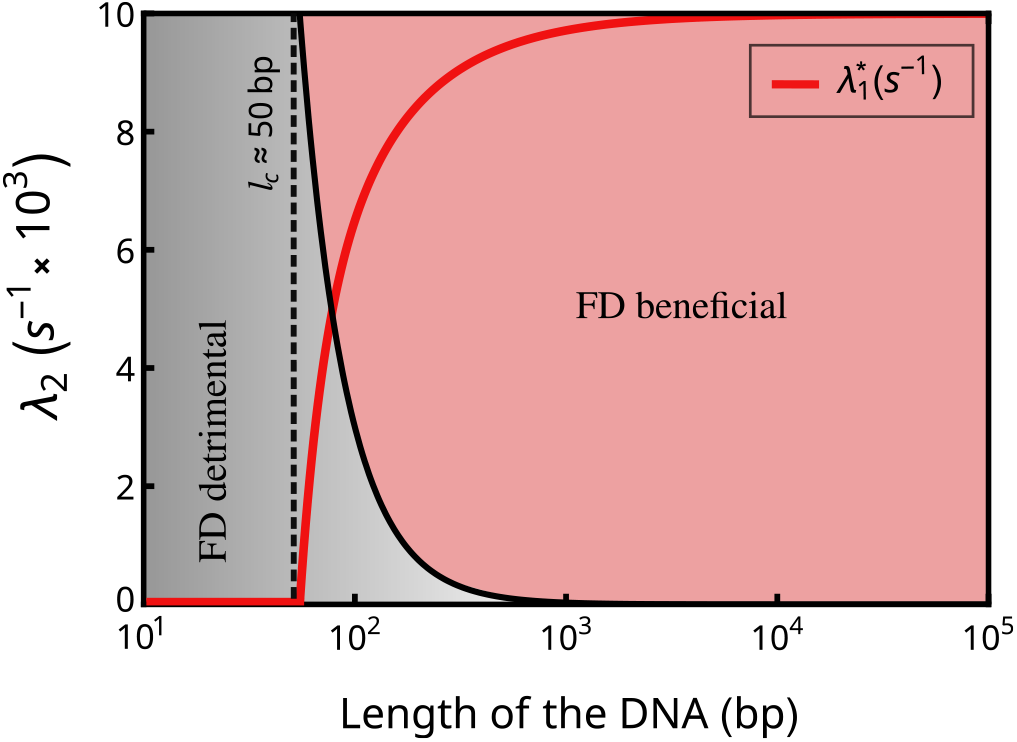
Phase diagram generated from the fluctuation inequality (11) in the *λ*_2_–length of the DNA parameter space. The diagram showcases two distinct regimes: one in which facilitated diffusion enhances the search efficiency, and another in which it fails to provide any acceleration. The boundary separating these regions—the separatrix(black curve)—is determined by the equality *CV*^2^ = 1 + 2 ⟨*T*_3D_ ⟩/ ⟨*T*_S_ ⟩. For all results shown here, the diffusion constant is fixed at 5 × 10^5^ bp^2^/s, as measured in case of E. coli bacteria for *lac-repressor* protein.^16^ The red curve corresponds to optimal detachment rate 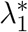 and the vertical dashed line indicates the critical length. This is plotted on the same axis as *λ*_2_ for different DNA lengths. In this case, the value of *λ*_2_ is fixed at 10^4^ s^−1^ as reported in.^30,67^

### Reduced fluctuations due to detachment

We now turn our attention to examine the fluctuations in the total search time for the facilitated diffusion mechanism. As shown in Fig. 5(a), the mean search time exhibits an optimal behavior when protein detachment from the DNA is introduced—indicating that intermittent unbinding can accelerate the search process. A natural question that follows is whether this detachment can also minimize the fluctuations in the total search time, in addition to reducing the mean. To address this, we derive an explicit analytical expression for the fluctuation, quantified by the standard deviation *σ*_Tot_. Through a detailed analysis of Eq. (17), we arrive at a compact and physically insightful expression for the variance under facilitated diffusion, which is presented below. The full derivation is provided in appendix -F. For symmetric DNA, we obtain the following expression

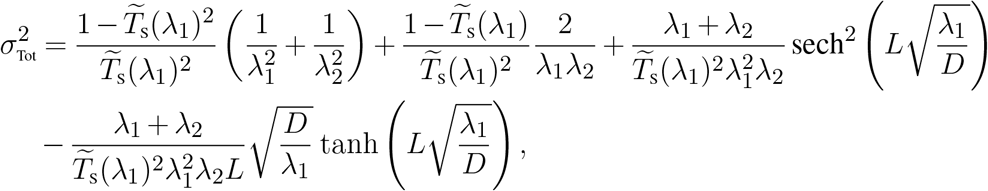

Where 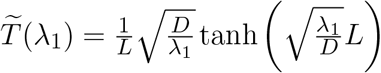. In Fig. 5(b), we plot the standard deviation of the total search time, *σ*_Tot_, as a function of the detachment rate *λ*_1_. The plot clearly reveals that the fluctuations in the search time can be reduced due to the detachment. This result highlights a remarkable dual role of detachment in the facilitated diffusion mechanism: while detachment helps expedite the mean search process by enabling the protein to escape local traps and explore new regions of the DNA, it simultaneously reduces variability in the overall search time. In other words, the same mechanism that enhances efficiency also controls variability.

## Conclusion

The problem of how proteins locate specific target sites on DNA is central to understanding numerous biological processes, including transcription, replication, genome regulation, and gene editing. This target search is inherently stochastic, shaped by the interplay between rapid local exploration and long-range relocations. Facilitated diffusion, wherein a protein alternates between one-dimensional (1D) sliding along the DNA and three-dimensional (3D) excursions through the surrounding medium, has long been recognized as a highly efficient mechanism for scanning extended genomic regions. However, despite extensive studies, a unifying principle governing when and why such combined dynamics outperform pure 1D or 3D search has remained elusive.

In this work, we employed a renewal-based first-passage formalism to analyze the facilitated diffusion process without imposing any specific assumptions on the microscopic details of either sliding or excursions. This approach led to three central results: a general expression for the mean and fluctuation of the total search time, how a combination of such intermittent dynamics can reduce both mean and fluctuations of the search time and a fluctuation inequality—expressed through a universal coefficient of variation (*CV*) relation—that dictates when intermittent detachment can reduce the mean search time. The *CV -criterion* reveals that the efficiency of the search critically depends on the balance between 1D sliding and 3D excursions. While detachment accelerates exploration on sufficiently long DNA, it becomes detrimental for short sequences; and even for long DNA, efficiency can break down when the mean excursion time ⟨*T*_3D_⟩ becomes large, as often occurs under macromolecular crowding,^8^ anomalous diffusion, or trapping in the intracellular environment.

The *CV -criterion* also uncovers a deeper physical principle underlying efficient target search: when 3D excursions are slow but not heavy tailed, the search can still remain efficient only if the 1D sliding process exhibits sufficiently large fluctuations. Broad distributions of sliding times, reflected in a high *CV*, naturally arise from biological features such as heterogeneous energy landscapes, deep kinetic traps, variable nonspecific binding affinities,^29–31^ obstacles from other DNA-bound proteins,^5,42–44^ and structural irregularities like loops^86^ or bends that locally confine the protein. These sources of stochasticity can broaden the 1D sliding time density, thereby ensuring that the *CV -criterion* can be satisfied even when ⟨*T*_3D_⟩ is large. In this sense, the universal *CV -criterion* provides a unifying explanation for when facilitated diffusion offers a true advantage, highlighting that the interplay between mean search time and fluctuations—not just their averages—governs optimal target search on DNA.

Finally, it is important to note that length is not the only parameter emerging as a signature of an efficient search as we have shown within our set-up. There may be additional parameters or DNA-specific properties—such as binding energy, curvature, or other structural features, that were not included in our model but could contribute to a more efficient facilitated diffusion mechanism. Earlier experiments have also reported subdiffusive motion of proteins along the DNA contour.^63^ Our framework is particularly well suited to quantify the role of detachment in such anomalous search dynamics and to examine the relevance of the *CV -criterion* in this context. Another promising extension is to incorporate hopping-mediated 3D excursions, which would render the excursion times spatially dependent.^63^ Our formalism naturally accommodates such generalizations and provides a systematic route to compute both the mean search time and its fluctuations in these realistic scenarios. Future works would explore these directions within this unified description of target search process.

## Author Contributions

Arnab Pal conceived and designed the research. Jitin Rajoria carried out the theoretical computations, numerical simulations, and analyzed the data. Jitin Rajoria and Arnab Pal wrote the article together.

## Acknowledgments

We thank Srabanti Chaudhury for fruitful discussion during the initial stage of this work. The numerical calculations reported in this work were carried out on the Nandadevi and Kamet cluster, which are maintained and supported by the Institute of Mathematical Science’s High-Performance Computing Center. We gratefully acknowledge research support from the Department of Atomic Energy, Government of India via Soft Matter Apex projects.

## Appendices

### A Derivation of the fluctuation inequality or the *CV -criterion*

In this section, we provide a derivation for the fluctuation inequality (11), as follows

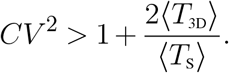

To this end, we begin with the inequality as discussed in the main text in Eq. (10)

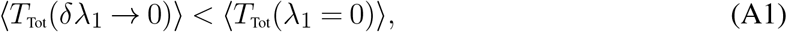

where ⟨*T*_Tot_(*λ*_1_ = 0)⟩ is the total search time without any detachment and therefore, it equates to ⟨*T*_S_⟩. To compute ⟨*T*_Tot_(*δλ*_1_ → 0)⟩, which is the average total search time when a very small detachment rate *δλ* is introduced, we do a linear response expansion of Eq. (7) to find

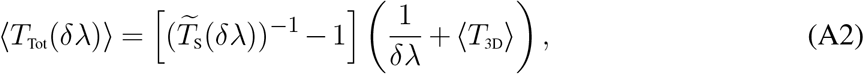

where 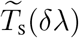 is the Laplace transform of the sliding time distribution 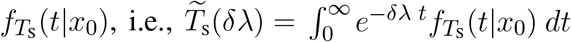, which can be expanded again using a Taylor’s series to find

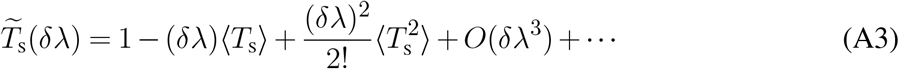

Substituting the above expansion into Eq. (A2) leads to the following expression for the mean time in the presence of an infinitesimal detachment rate

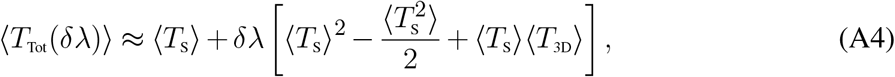

where we have neglected the higher-order term, i.e., *O*(*δλ*^2^) and so on. Substituting the above expression into the inequality (A1), we find

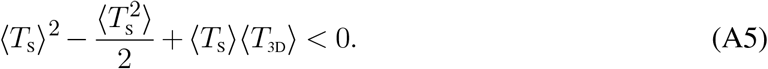

After a simple rearrangement of terms, we can write

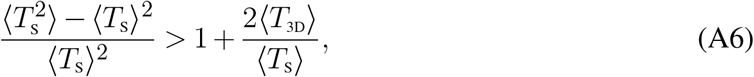

where the quantity on the left hand side can be identified as *CV*^2^ so that we arrive at inequality (11). This concludes the derivation.

### B Derivation of the backward Fokker-Planck equation for 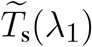 – Eq. (21)

In this section we provide a proof for Eq. (21). To this end, we first define *Q*(*t*|*x*_0_), which is known as the survival probability i.e., the probability that the protein has not been able to detect the target site upto time *t*, given that it had started from a fixed position *x*_0_ (to be averaged over at the end) along the DNA. By definition,

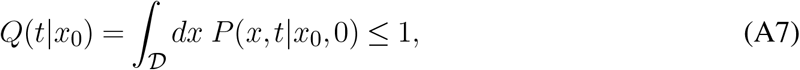

where the domain of search is indicated by 𝒟 and furthermore, it can be shown to satisfy a diffusion-alike (recall Eq. (18)) backward Fokker-Planck equation given by^74,84^

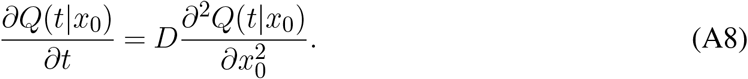

Eq. (A8) needs to be supplemented with one initial condition *Q*(*t*|*x*_0_)|_*t*=0_ = 1 and two boundary conditions. There are two regions in which the above differential equation (A8) has to be solved separately: −*L < x*_0_ *<* 0 and 0 *< x*_0_ *< R*. If the protein is located at position *x*_0_ = 0, then the detection is immediate and thus, one of the boundary conditions is 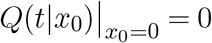. Since there are reflecting boundaries at −*L* and *R*, the probability flux to these two coordinates should be zero there i.e., 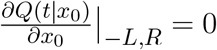.^74,84^ The survival probability *Q*(*t*|*x*_0_) and sliding time density 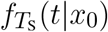 are related to each other by^74,84^

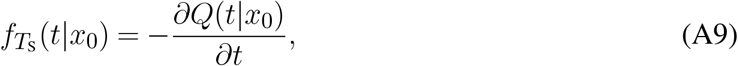

so that Eq. (A8) simply transforms to an ordinary differential equation

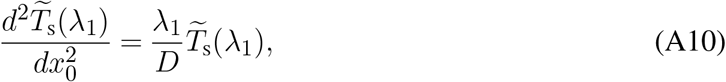

where recall 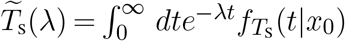 is the Laplace transform of the sliding time density function. Note that the transformed boundary conditions now read 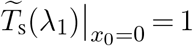 and 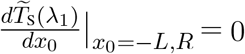. Solving Eq. (A10) systematically within the regions with the above-mentioned boundary conditions and finally, performing an average over the uniformly distributed initial positions, we arrive at Eq. (22).

### C Derivation of distribution function for the total search time

In model section, we have provided a compact expression for the moment generating function given by Eq. (15). In this section, we provide a detailed derivation of the same. We start by recalling Eq. (3)

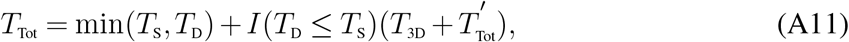

which can be rewritten in the following way

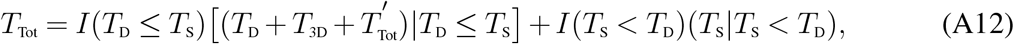

where recall the definition of an indicator random variable *I*(*z*_1_ ≤ *z*_2_) = 1 when *z*_1_ ≤ *z*_2_, and zero, otherwise. Performing an average over both the random variables *z*_1_, *z*_2_ yields ⟨*I*(*z*_1_ ≤ *z*_2_)⟩ = Prob(*z*_1_ ≤ *z*_2_), this property will be used later in the derivation. Let us now define the distribution function of the total search time in Laplace space, often denoted as the moment generating function (MGF), in the following way

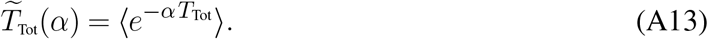

Substituting Eq. (A12), we can rewrite the MGF as

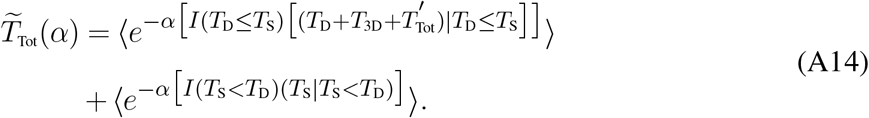

Taking the total expectation and using the property of the indicator functions, we can rewrite the above expression in the following way

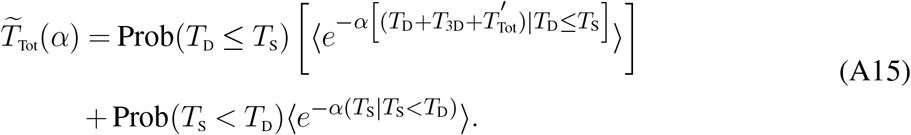

Before proceeding further, it will be useful to recall the conditional times as defined in Eq. (12) of the main text

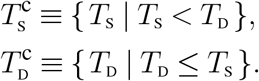

Using these definitions and further utilizing the property that 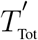 is an identical & independent copy of *T*_Tot_ from a different trial so that 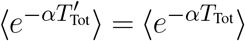 and consequently, is independent of 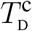 and *T*_3D_, we rewrite Eq. (A15) in the following way

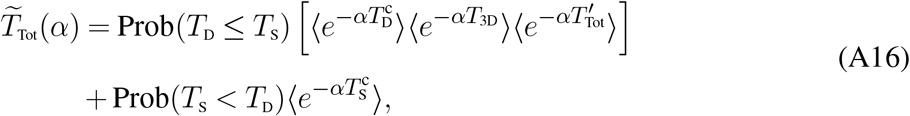

Recalling Prob(*T*_S_ *< T*_D_) = *q* from Eq. (2), together with the definitions of the Laplace-space distribution functions for the various times, we arrive at the following expression

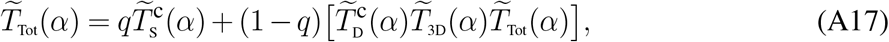

which can be rearranged to finally obtain

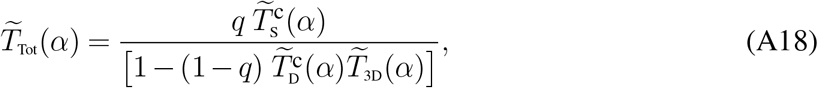

which was announced in the model section of the main text as in Eq. (15). This expression for the MGF is extremely useful to compute the mean and higher order of moments in search time. In addition, it can also provide insights to the large time asymptotics for the search time density by doing a small *α* → 0 expansion.

### D Derivation for fluctuations in terms of standard deviation

This appendix provides important steps to calculate an expression for the standard deviation of the total search time *T*_Tot_. The standard deviation of the total search time is given by

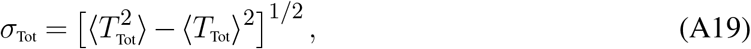

which requires formal expressions of the first and second moment. In general, *n*-th order moment can be extracted from

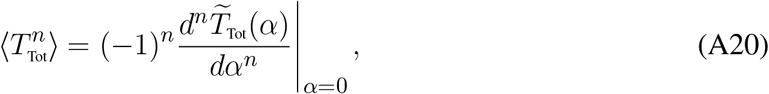

where the MGF, given by Eq (A18), has been derived in the last section. Differentiating Eq. (A18) once and taking *α*→0, with the identification that

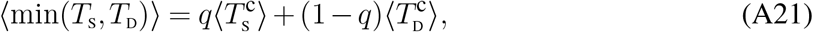

we recover,

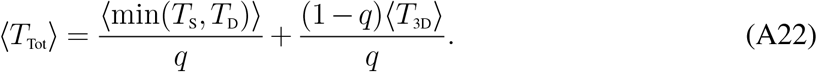

Now, we differentiate twice and and take *α*→0, with the identification that 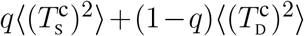 is equivalent to ⟨min(*T*_S_, *T*_D_)^2^⟩ so that

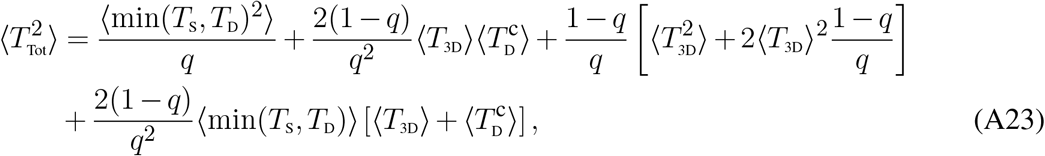

which was announced in the main text. So far, most of these expressions remain formal. Although they are useful for theoretical computation, for practical purposes, we need to connect those with the various time-densities which are experimentally measurable observables. This is done in the next Appendix.

### E Connecting the average quantities with the time-densities

In here, we show how to connect the observables with the accessible time-densities. For instance, *q* = Prob(*T*_S_ *< T*_D_) can be written either of the following way based on the order of conditioning for the random times

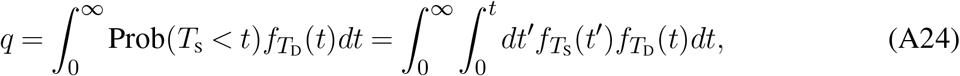

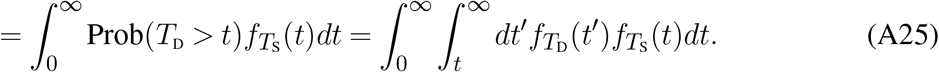

To calculate the average of conditional detachment time 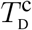, we use the probability distribution given in Eq. (14). The expression for the average can therefore be written in the following way

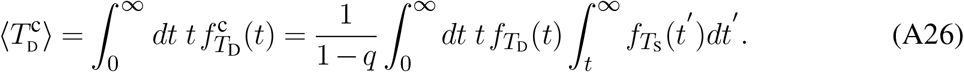

Extraction of the higher moments also follows the same procedure. A similar expression can be written for the conditional time 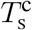 as well, given by

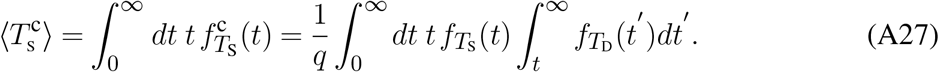

Our next goal is to compute the quantities associated with the minimum of the two times *T*_S_ and *T*_D_. To find the average of min(*T*_S_, *T*_D_) and min(*T*_S_, *T*_D_)^2^, we first evaluate the probability distribution of minimum. We utilize the concept of probability theory to find the distribution of a random variable. The distribution is given below

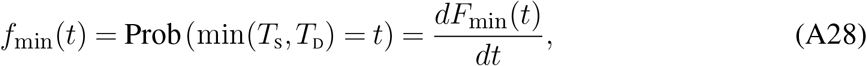

where *F*_min_(*t*) is the cumulative distribution function which is defined in the following way

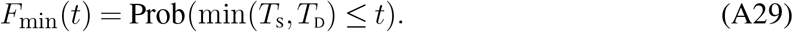

This can be evaluated using the following steps assuming *T*_S_ and *T*_D_ are independent so that

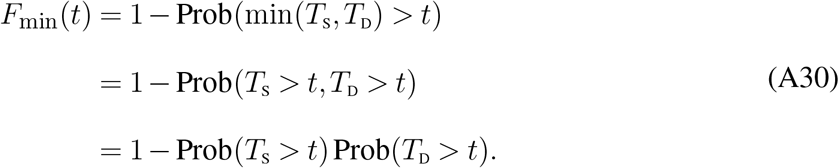

Consequently, the probability distribution function attains the following form

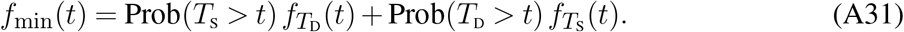

Finally, any moment of min(*T*_S_, *T*_D_) can be evaluated by using this probability distribution function. The general form is given below

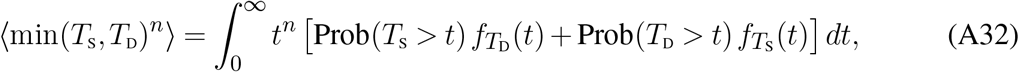

where 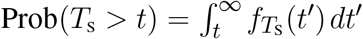 and 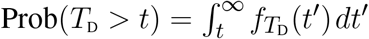. All four quantities are calculated for arbitrary time density functions 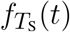 and 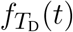.

### F The specific form of probabilities and averages for the exponentially distributed detachments

We now consider the case where the detachment time is assumed to follow an exponential distribution, i.e., 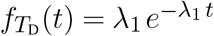. In that case, the observables reduce to much simpler forms. Namely, the success probability *q* becomes the Laplace transform of the successful sliding time distribution

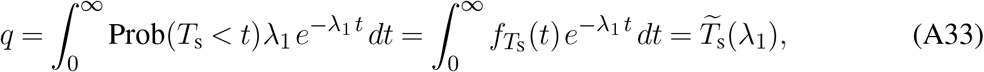

where we have used the definition of the probability distribution in terms of CDF from Eq. (A28). The average conditional detachment time, following Eqs. (A26) and (14), can be rewritten in the following way

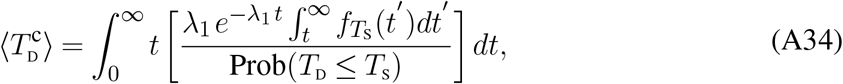

where the denominator is simply 1 − *q*, as discussed previously. Therefore, 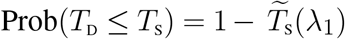. We perform integration by parts and use 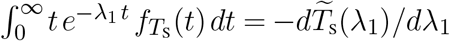 along with 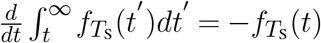 so that

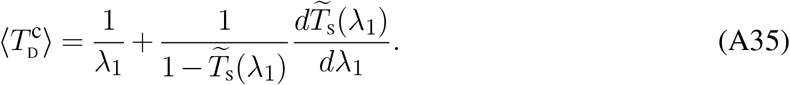

Similarly, we find the mean conditional sliding time to be

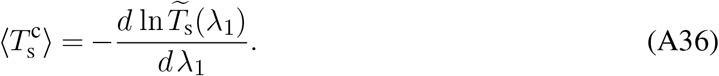

Substituting these expressions in Eq. (A32), we find

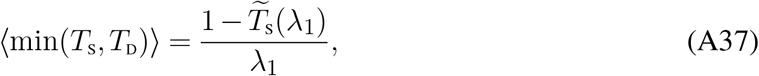

and

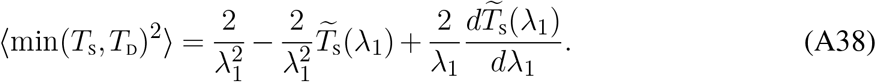

Further assuming the 3D excursion events to be rate processes in the case of facilitated diffusion, we find

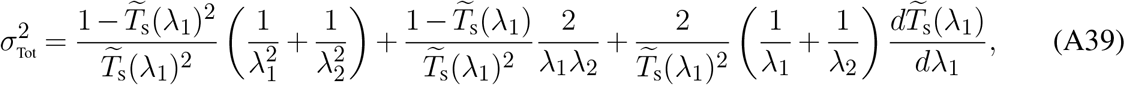

where we have assumed the mean time in a 3D excursion to be 1*/λ*_2_, and 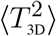 is 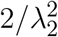. To recover the final result in Eq. (27), we recall Eq. (22) which reads

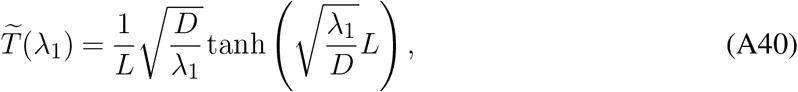

substituting which in Eq. (A39), we arrive at Eq. (27)

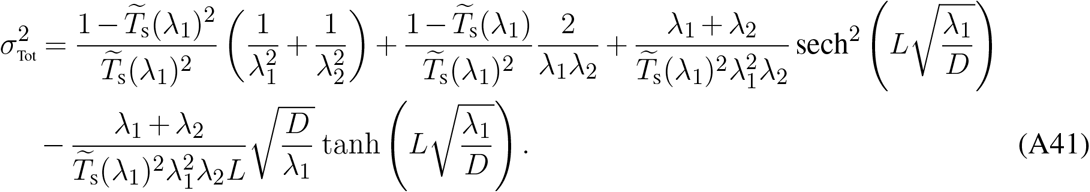

